# Early alterations in cortical and cerebellar regional brain growth in Down Syndrome: An in-vivo fetal and neonatal MRI assessment

**DOI:** 10.1101/683656

**Authors:** Prachi A. Patkee, Ana A. Baburamani, Vanessa Kyriakopoulou, Alice Davidson, Elhaam Avini, Ralica Dimitrova, Joanna Allsop, Emer Hughes, Johanna Kangas, Grainne McAlonan, Mary A. Rutherford

## Abstract

Down Syndrome (DS) is the most frequent genetic cause of intellectual disability with a wide spectrum of neurodevelopmental outcomes. At present, the relationship between structural brain morphology and the spectrum of cognitive phenotypes in DS, is not well understood. This study aimed to quantify development of the fetal and neonatal brain in DS using dedicated, optimised and motion-corrected *in-vivo* magnetic resonance imaging (MRI). We detected deviations in development and altered regional brain growth in the fetus with DS from 21 weeks’ gestation, when compared to age-matched controls. Reduced cerebellar volume was apparent in the second trimester with significant alteration in cortical growth becoming evident during the third trimester. Developmental abnormalities in the cortex and cerebellum are likely substrates for later neurocognitive impairment, and ongoing studies will allow us to confirm the role of antenatal MRI as an early biomarker for subsequent cognitive ability in DS. In the era of rapidly developing technologies, it is hoped that the results of this study will assist counselling for prospective parents.

## Introduction

Down Syndrome (DS) is the most common genetic cause of intellectual disability, occurring as a result of triplication of a genomic region on human chromosome 21 (Trisomy 21; T21) ^1,2^. The neurodevelopmental phenotype of DS is associated with cognitive deficits with varying degrees of impairment of speech, motor and language functions, with a wide and largely unexplained spectrum of IQ’s (range = 20-80; mean = 50) ^3–6^. Approximately 50% of DS babies are born with a congenital heart defect (CHD), the most common of which are endocardial cushion defects, including atrioventricular septal defects (AVSD; 45%) and ventricular septal defects (VSD; 35%) ^7–10^. The presence of cardiac disease is associated with a poorer outcome, as observed in typically developing children with CHD, where studies have shown deficits in neurocognitive, motor and psychosocial skills ^11^. Children with DS who have an AVSD had poorer gross motor skills, cognition and lower scores in expressive and receptive vocabulary, as compared to DS children with a structurally normal heart ^12–14^. However, there is limited information regarding structural brain development in the presence or absence of CHD in DS.

The severity and range of problems experienced in DS are determined, in part, by overexpression of specific genes on the extra chromosome, and dysregulation of their associated pathways ^15,16^. At present, there is little known about early brain growth trajectories in DS and the spectrum of cognitive phenotypes. Abnormalities in brain development in DS, such as reductions in cortex and cerebellum have been observed in early life, predominantly through post-mortem cases, but there is little *in vivo* information about when atypical brain growth and development patterns may emerge ^17^. Ultrasound studies have concentrated on first trimester findings, to assist in early diagnosis ^18^, with a few small studies focusing on the second and third trimester, and noting smaller head circumferences and cerebellar diameters ^19^.

Previous studies have largely focused on children and adults with DS, which makes it difficult to untangle the primary pathophysiological and direct effects of the condition on the brain, from the secondary or compensatory mechanisms which arise from living with the disorder; additionally, assessing the impact of indirect effects of cardiac compromise on postnatal brain development. To reduce these confounds, a more detailed understanding of early *in vivo* growth of the cortex and cerebellum, may allow us to identify early biomarkers to predict the severity of initial cognitive impairments, and improve counselling for prospective parents. It also has the potential to reveal new windows for future therapeutic intervention strategies aimed at improving neurodevelopmental outcomes.

The primary objective of this study was to investigate the developing brain across gestation, in a cohort of fetuses and neonates with DS, with and without an associated cardiac defect, using MR imaging techniques. We aimed to objectively quantify three-dimensional (3D) volumetric and two-dimensional (2D) linear measurements of whole brain, cortex and cerebellum and to compare these parameters with brain measures from aged-matched control cases. We hypothesised that there are quantifiable alterations in early cortical and cerebellar growth in fetuses and neonates with DS as compared with controls and that these are further altered by the presence of a cardiac abnormality.

## Methods

Ethical approval for this study was obtained from the West London and GTAC Research Ethics Committee (REC) for DS participants and fetal controls (07/H0707/105); and from the Dulwich NREC for neonatal controls (12/LO/2017). Informed consent was obtained from all participants or their legal guardians, prior to imaging at Hammersmith Hospital and St. Thomas’ Hospital, London, UK. All methods were carried out in accordance with the relevant guidelines and regulations.

### Participants

Pregnant women with a fetus diagnosed with DS or with a high risk for DS were referred from antenatal and fetal medicine clinics across London hospitals, following their anomaly scans, amniocentesis, or blood test results. Ex-fetal participants who consented during their initial scan to be contacted post-delivery, were invited for a neonatal scan up to 46 weeks PMA (post-menstrual age). Additionally, neonates with confirmed DS were also recruited from the neonatal unit or postnatal wards at St Thomas’ Hospital. Participants with DS and other non-brain congenital abnormalities such as cardiac defects or gastrointestinal malformations were also included in this study. The presence and characteristics of any antenatally detected congenital heart defect were confirmed on echocardiography postnatally as part of routine clinical assessment. Clinical details for all fetuses and neonates with DS can be found in supplementary materials (Table 4).

Healthy pregnant volunteers were recruited from the antenatal clinics at Queen Charlotte’s and Chelsea Hospital and St. Thomas’ Hospital, London at the time of their anomaly scans (approximately 20 weeks’ gestational age) or self-referred. The entry criterion for a low risk control for this study, was a normal brain appearance as determined by fetal MRI and confirmed by an experienced perinatal neuroradiologist. Fetuses with other congenital or chromosomal abnormalities, women with a history of drug use, twin pregnancies, pregnancies with delivery complications resulting in neonatal neurological complications were excluded from the study. In addition, participants with an abnormal appearance of the brain on neonatal MRI, birth weight below the 3rd centile, premature birth (<37 weeks GA), sub-optimal MR image quality and with an abnormal outcome at either the year 1 or 2 neurodevelopmental assessments were also excluded. Control group neonates were recruited from South London and South East of England antenatal centres as part of the Brain Imaging in Babies Study (BIBS), and scanned at the Centre for the Developing Brain (St. Thomas’ Hospital, London).

### Fetal Image Acquisition

Fetal MRI was performed on a 1.5-Tesla or 3-Tesla MRI System (Philips Achieva; Philips Medical Systems, Best, The Netherlands) using a 32-channel cardiac array coil placed around the maternal abdomen. Participants were scanned in either left lateral or supine position and provided with a pregnancy pillow and foam wedges for comfort. Maternal temperature was taken before and after the scan, and heart rate was monitored throughout. Sedatives were not administered and the total scan time was restricted to 60 minutes.

Fetal T2-weighted images were acquired in the transverse, sagittal and coronal plane using single shot turbo spin echo (ssTSE) sequences with the following scanning parameters: repetition time (TR) = 15 s; echo time (TE) = 160 ms; slice thickness = 2.5 mm with a slice overlap = 1.5 mm; and flip angle = 90 degrees. brain was oversampled by acquiring multiple overlapping single shot T2-weighted images to ensure complete coverage of all regions of interest within the brain.

3D reconstructed images were obtained using Snapshot MRI with Volume Reconstruction (SVR) from 2D slices ^20,21^. A stack of images with the least motion and artefacts was selected as the target, and a mask was created to isolate the fetal brain from surrounding maternal and extracranial fetal tissue, using ITK-SNAP (version 2.4.0; Yushkevich et al. 2006). The optimised reconstruction algorithm used intensity matching to exclude mis-registered or corrupted voxels and slices. The approved slices were registered into a common coordinate space for alignment, to produce the resultant high signal-to-noise ratio images. The reconstructed brains were manually re-orientated and then resampled to a voxel size of 0.2 × 0.2 × 1 mm, to improve the resolution of smaller structures. The data sets provide complete coverage of the brain in the orthogonal planes, which is optimum for detailed volumetric analysis.

### Neonatal Image Acquisition

Neonatal MR scanning was performed on a Philips Achieva 3-Tesla system (Best, The Netherlands) at the Centre for the Developing Brain (St. Thomas’ Hospital, London) using a dedicated 32 channel neonatal head coil. Infants were placed and secured within the scanning shell and a neonatal positioning device was used to stabilise the head to reduce movement ^23^. Auditory protection was comprised of earplugs moulded from silicone-based putty placed in the outer ear (President Putty, Coltene/Whaledent Inc., OH, USA), neonatal earmuffs over the ear (MiniMuffs, Natus Medical Inc., CA, USA) and an acoustic hood placed over the scanning shell. Sedation was not administered and all babies were scanned during natural sleep. An experienced neonatologist was present during the examinations and pulse oximetry, heart rate, and temperature were monitored throughout the scan.

T2-weighted images were acquired in the sagittal and transverse planes using a multi-slice turbo spin echo sequence. Two stacks of 2D slices were acquired using the scanning parameters: TR = 12 s; TE = 156 ms; slice thickness = 1.6 mm with a slice overlap = 0.8 mm; flip angle = 90 degrees and an in-plane resolution: 0.8×0.8 mm.

Visual analysis of all fetal and neonatal images was performed by a specialist perinatal radiologist to exclude additional anomalies and confirm appropriate appearance for gestation. Image datasets showing an overt additional malformation were excluded. All T2-weighted images were reviewed to assess image quality and those with excessive motion, which could not be automatically accurately segmented, were excluded.

### Image Analysis

#### Linear Measurements

Fetal and neonatal brain linear measurements were performed on reconstructed images using ImageJ (version 1.46R, National Institutes of Health, Bethesda, MD, USA), and based on the following parameters ^24,25^. Fig. 4; (a) Biparietal diameter (BPD) brain – maximum brain width in the transverse plane, outermost to outermost edge; (b) Biparietal diameter (BPD) skull – the widest diameter of the fetal skull measured in the transverse plane, outermost to innermost edge; (c) Occipitofrontal diameter (OFD) brain – the maximum distance of the brain width between the frontal and occipital lobes, measured in the sagittal plane, outermost to outermost edge; (d) Occipitofrontal diameter (OFD) skull the maximum distance between the frontal and occipital skull bones, measured in the sagittal plane, outermost to innermost edge (e) Cavum width - largest width in the transverse plane; (f) Head circumference (HC) – calculated using the equation: head circumference = 1.62 × [(skull BPD) + (skull OFD)]; (g) Transcerebellar diameter (TCD) – the maximum lateral cerebellar distance in the transverse plane. All vermis measurements were taken in the mid-sagittal plane, (h) Vermis height – the maximum superior-inferior length; (i) Vermis width – the maximum distance between the fastigium and the posterior part of the vermis; and (j) Vermis area – calculated using a free hand drawing tool.

**Figure 1:**
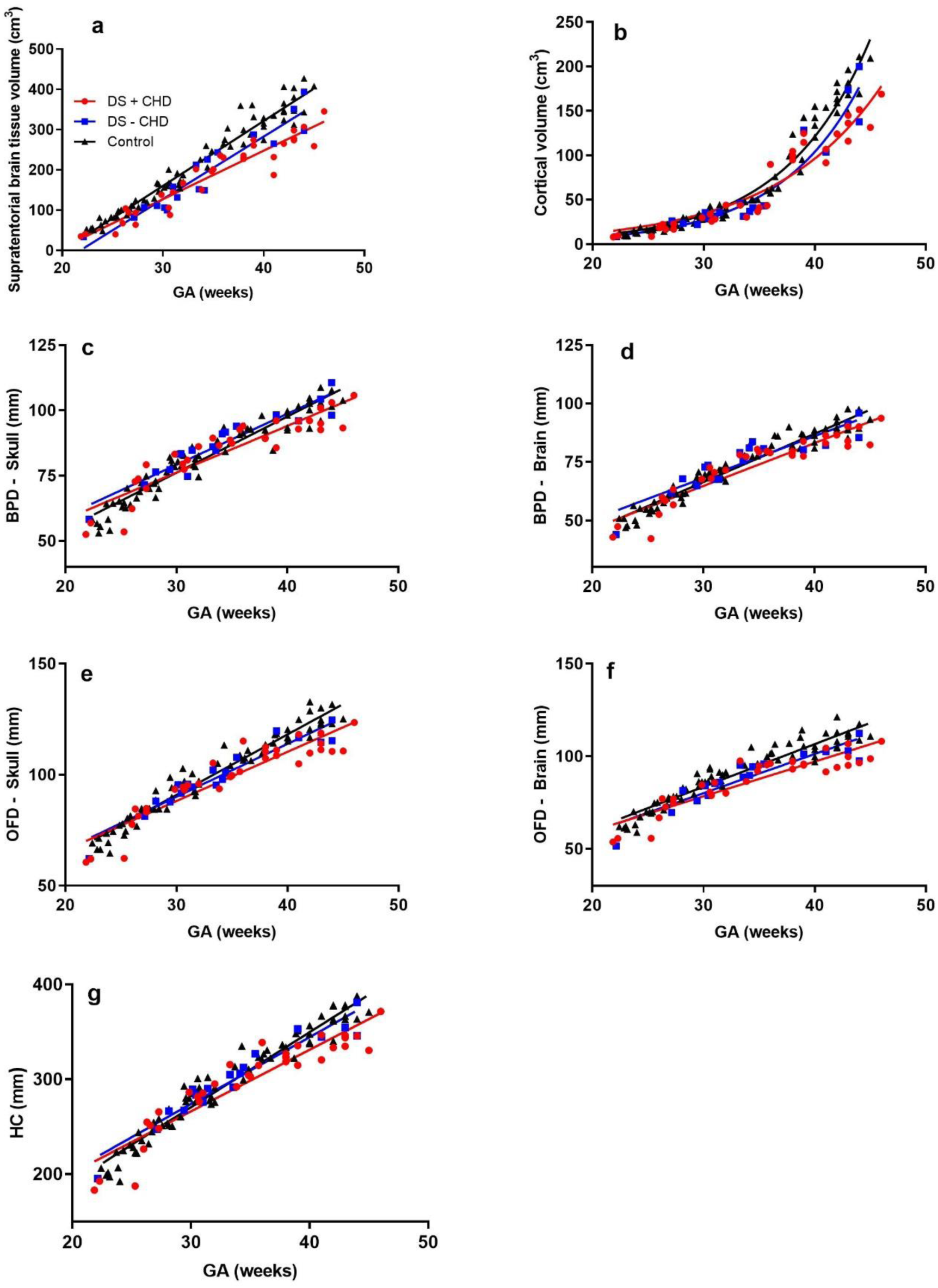
2D and 3D regional measures of the brain in fetuses and neonates with DS and a congenital heart defect (DS+CHD; red circles), DS without a congenital heart defect (DS–CHD; Blue squares), and age-matched normal controls (black triangles). (a) Whole brain volumes, (b) Cortical volumes, (c) Biparietal diameter (BPD) - skull, (d) Biparietal diameter (BPD) - brain, (e) Occipitofrontal diameter (OFD) – skull, (f) Occipitofrontal diameter (OFD) - brain, and (g) Head circumference (HC) (MRI derived).

**Figure 2:**
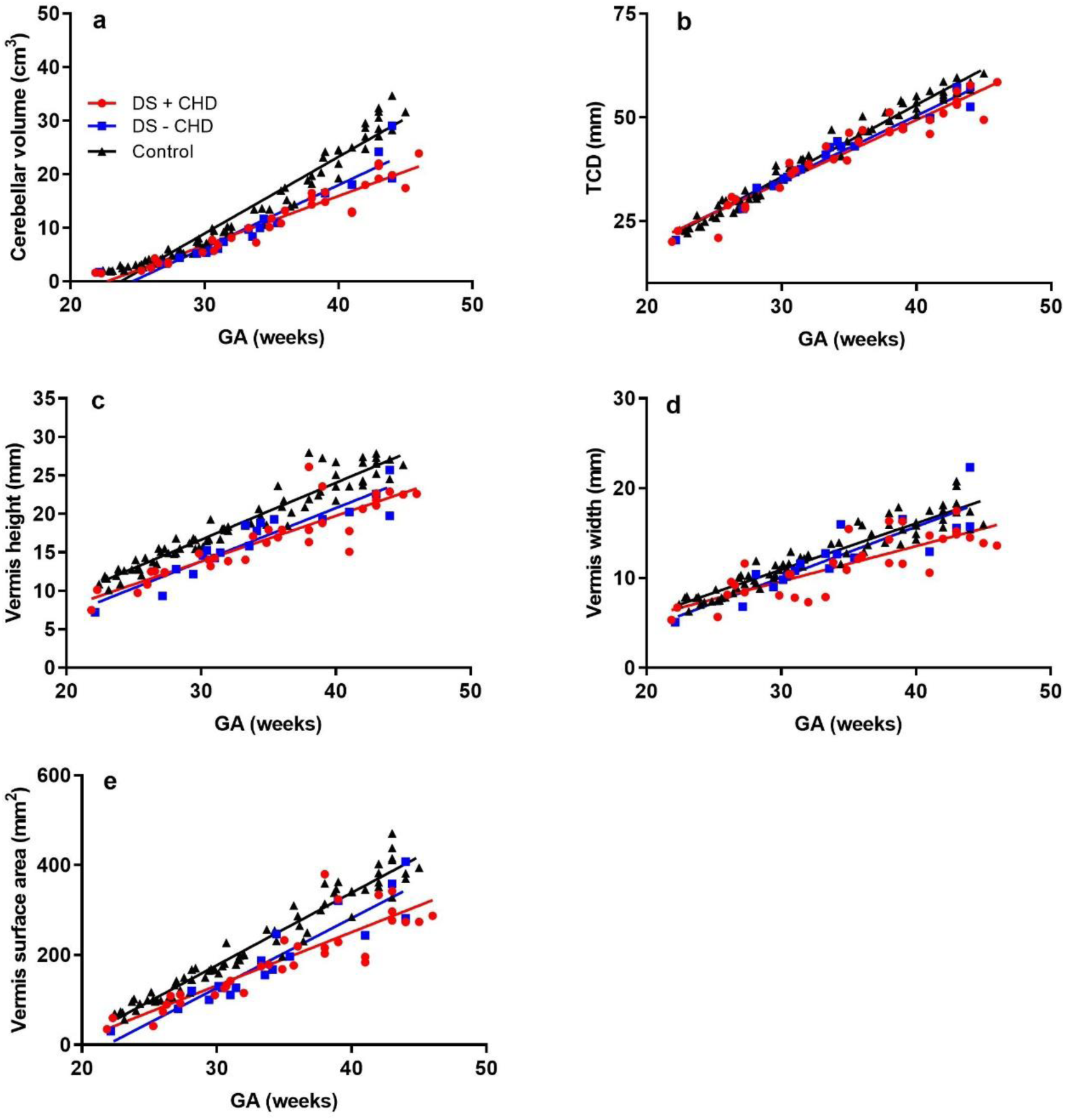
2D and 3D measures of the cerebellum in fetuses and neonates with DS and a congenital heart defect (DS+CHD; red circles), DS without a congenital heart defect (DS–CHD; blue squares), and age-matched normal controls (black triangles). (a) Cerebellar volume, (b) Transcerebellar diameter, (c) Vermis height (d) Vermis width, and (e) Vermis surface area.

**Figure 3:**
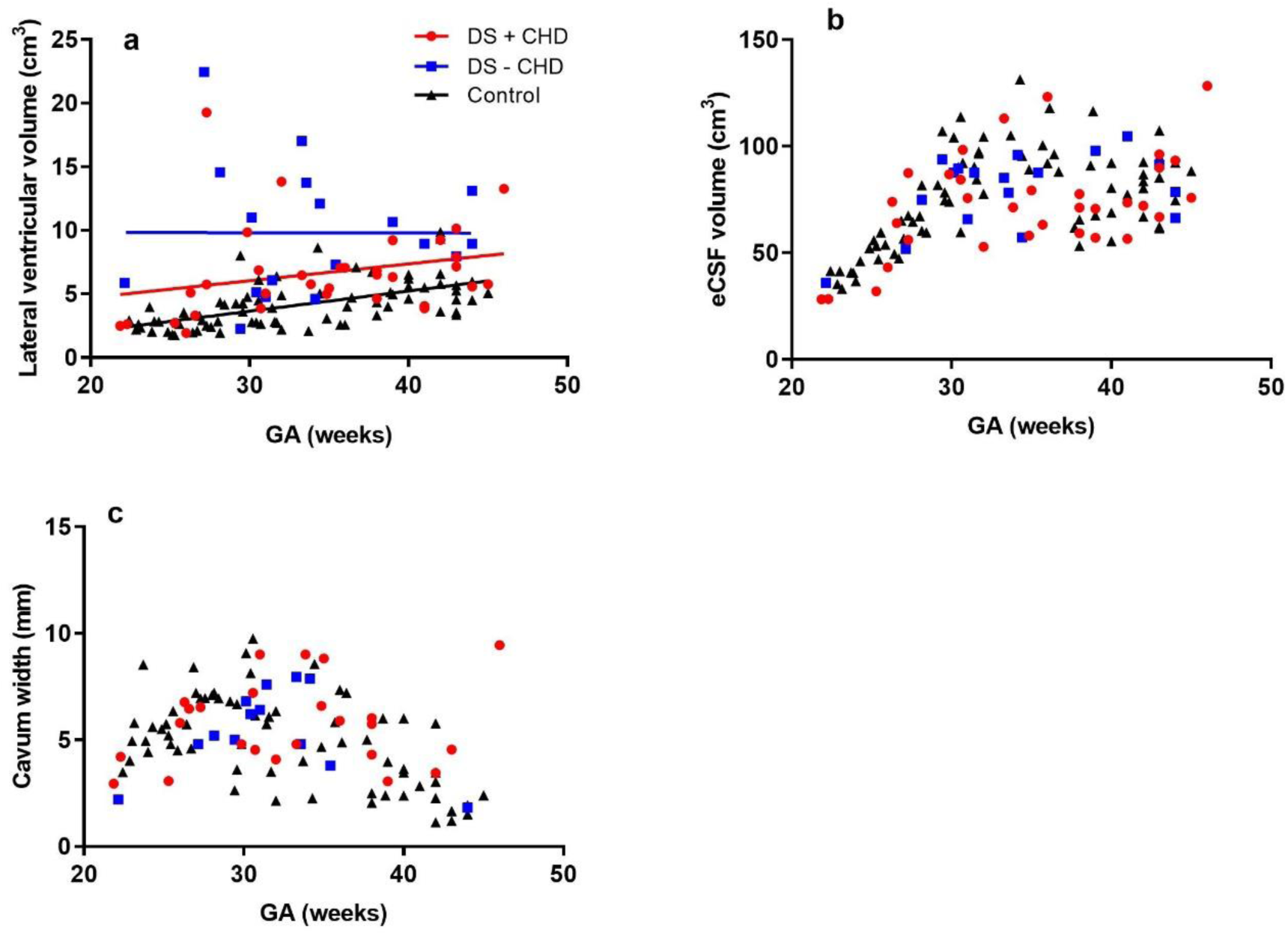
2D and 3D measures of the CSF regions in fetuses and neonates with DS and a congenital heart defect (DS + CHD; red circles), DS without a congenital heart defect (DS – CHD; blue squares), and age-matched normal controls (black triangles). (a) Lateral ventricular volume, (b) eCSF volume, and (c) Cavum width.

**Figure 4:**
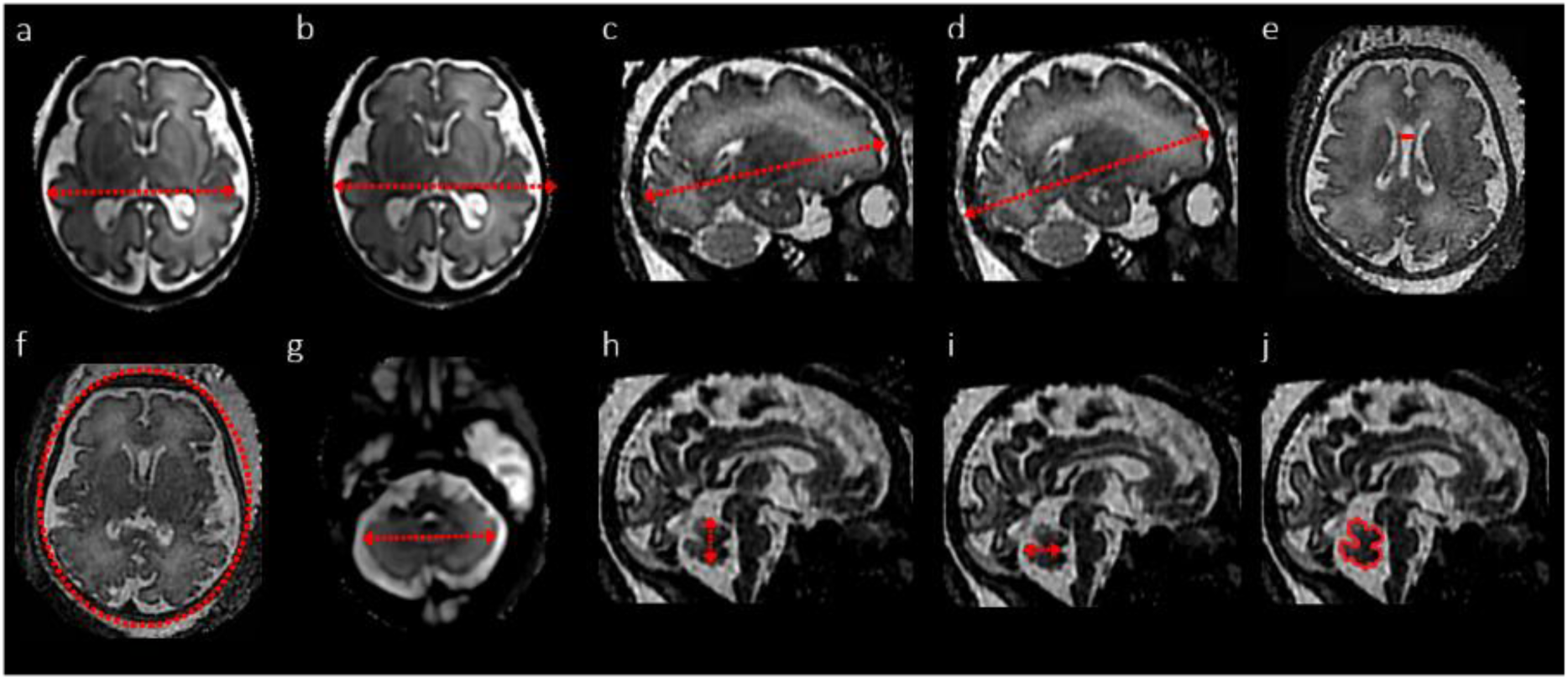
Visual representation of linear measurements in the fetal brain (a: BPD – brain, b: BPD – skull, c: OFD brain, d: OFD – skull, e: cavum width, f: HC, g: TCD; h: vermis height; i: vermis width; j: vermis surface area).

#### Fetal volumetric segmentation

Volumetric segmentation of the supratentorial brain tissue, lateral ventricles and extra-cerebral CSF (eCSF) was performed in a semi-automated manner, whilst the cortex and cerebellum were manually segmented using ITK-SNAP ^22^ using the following parameters; (a) Supratentorial brain tissue volume - brain tissue above the tentorium, excluding the brainstem, cerebellum and any CSF spaces; (b) Cortical volume - total cerebral cortical grey matter; (c) Total ventricular volume - combined volumes of the left and right ventricles, including the choroid plexus, but excluding the third and fourth ventricle, the cavum septum pellucidum and cavum vergae (CSP); (d) eCSF volumes - all the intracranial CSF spaces surrounding the supratentorial brain tissue and cerebellum and including the interhemispheric fissure but not any ventricular structure, the CSP or vergae; and (e) Total cerebellar volume - both cerebellar hemispheres and the vermis and excluding the pons and fourth ventricle (Fig. 5a-e). To ensure compatibility between volumetric data acquired from 1.5T and 3T reconstructions, a healthy pregnant volunteer was scanned on both scanners on the same day. Volumetric data was compared and no significant differences were found between the two images across any of the key fetal brain segmentation regions; volumes were within the excepted +/-5% threshold.

**Figure 5:**
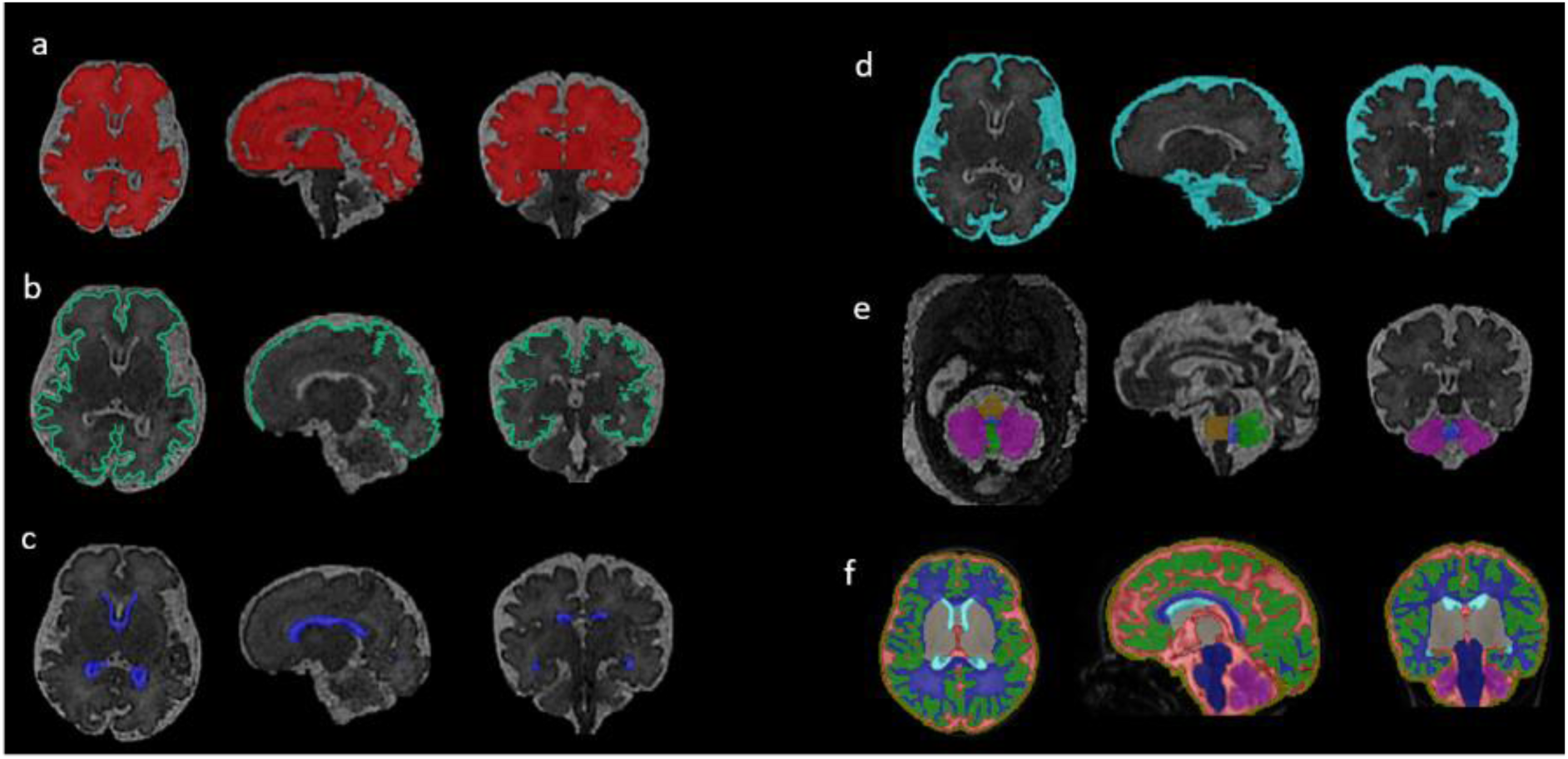
Volumetric segmentation of the fetal brain. Segmentation of T2-weighted volumetric MR images showing (a) supratentorial brain tissue (red), (b) cortex (green), (c) lateral ventricles (dark blue), (d) extra cerebral cerebrospinal fluid (light blue), (e) cerebellar hemispheres (pink), cerebellar vermis (bright green), pons (yellow) and fourth ventricle (blue); and (f) Neonatal T2 automatic atlas-based segmentation with 9 regions of interest.

#### Neonatal volumetric segmentation

T2-weighted images were segmented using *The Developing Brain Region Annotation With Expectation-Maximization (Draw-EM*) MIRTK Package, a fully automated tissue segmentation algorithm, optimised for the neonatal brain ^26^. Tissue segmentations were visually inspected for accuracy and any mislabelled voxels were manually edited using ITK-SNAP. Volumetric analysis was performed on the same five regions of interest as the fetal population. Supratentorial brain tissue volumes were calculated by the sum of cortical grey matter, deep grey matter and white matter volumes (Fig. 5f).

### Statistical analysis

Statistical analysis was performed using the SPSS software package (SPSS Chicago, IL, USA; version 24) and MATLAB (Release: 2017a, The MathWorks, Inc., Natick, MA, USA), and graphs produced on GraphPad Prism (GraphPad software, La Jolla, CA, USA; version 8.00). Each variable was tested for normality, non-normally distributed data was log-transformed. Linear and volumetric measurements for the combined fetal and neonatal cohorts were analysed using linear mixed-effects models (LME) to compare results between the two groups (DS and controls), controlling for the covariate GA/PMA as a fixed effect, and with a subject dependent random effect for the intercept to account for repeated measures. A post-hoc False Discovery Rate (FDR) test was performed for multiple comparisons, using the Benjamini-Hochberg (1995) procedure, controlling the alpha error to 5%. Correlations were assessed using the Pearson’s correlation coefficient (Pearson’s r) on normally distributed data and the Spearman’s correlation coefficient (Spearman’s r) on non-normally distributed data. The inter-rater and intra-rater variability was assessed for each region of interest and the threshold for differences was set at +/-5%.

## Results

### Control Cohort

A total of 52 control fetuses (mean gestational age (GA): 29.58 weeks; GA range of 22.43 – 38.86 weeks), and 21 control neonates (mean post-menstrual age (PMA): 41.58 weeks; PMA range: 38.00-46.00 weeks) were included as part of this study.

### DS cohort

30 fetuses with DS, of which 17 fetuses with cardiac defects (mean GA: 29.28 weeks; GA range: 21.86-35.71 weeks) and 13 without cardiac defects (mean GA: 30.82 weeks; GA range: 22.14-35.43 weeks) were included in the study. Similarly, 21 neonates with DS were scanned; 16 with cardiac defects (mean PMA: 40.51 weeks; PMA range: 36.00-46.00 weeks) and 5 without cardiac defects (mean PMA: 42.20; PMA range: 39.00-44.00). Additionally, 6 fetuses were scanned as neonates, and 8 neonates were born prematurely (GA at birth 31.00-37.00 weeks) (Table 1). The DS cohort was comprised of 22 female and 21 male participants (Supplementary Materials, Table 4).

**Table 1:** Combined fetal and neonatal DS and control cohorts, pre and post 28 weeks’ gestational age (GA); post-menstrual age (PMA)

|  | Control |  | DS |  |
| --- | --- | --- | --- | --- |
|  | GA < 28.00 weeks | GA > 28.00 weeks | GA < 28.00 weeks | GA > 28.00 weeks |
| Total number | N = 20 | N=53 | N = 10 | N = 41 |
| GA/PMA range (weeks) | 22.43-27.57 | 28.00-45.00 | 21.86-27.14 | 28.14-46.00 |
| Mean GA/PMA at scan (weeks) | 25.07 | 36.05 | 25.22 | 36.71 |

The first line of analysis aimed to compare regional brain volumes between the DS cohort and the controls during the second trimester (GA < 28.00 weeks) and third trimester into the neonatal period (GA > 28.00 weeks) (Table 1). Secondary analysis was performed to investigate the effect of a congenital cardiac defect on regional brain volumes in the DS cohort – cohorts were examined as a whole across gestation and postnatally and not subdivided by trimester (Table 2).

**Table 2:** Summary of the DS cohort, with and without a congenital heart defect (CHD)

|  | DS |  |
| --- | --- | --- |
|  | Cardiac abnormalities<br>(DS + CHD) | Non-cardiac<br>(DS – CHD) |
| Total number | N = 33 | N = 18 |
| GA/PMA range (weeks) | 21.86-46.00 | 22.14-44.00 |
| Mean GA/PMA at scan (weeks) | 34.72 | 33.98 |

### Whole brain and cortical development

All volumetric and linear brain and skull measures were positively correlated with increasing GA (Fig 1). All 3D brain measures increased with GA. The fetal and neonatal cohort with DS were found to have significantly smaller whole brain volumes in the second and third trimester and postnatally (supratentorial brain tissue (p < 0.01)). However, cortical volumes only started to deviate from the normal cohort in the third trimester (p < 0.01), after 28 weeks GA (Table 3, Fig. 1 (a, b). No differences in whole brain or cortical volumes were found between the DS fetuses and neonates with a congenital cardiac defect (DS + CHD) and those without (DS – CHD). Whole brain volumes were smaller in female participants with DS, along with smaller BPD skull and vermis width, however these were no longer significant when corrected for whole brain volume.

**Table 3:** Summary statistics for 2D and 3D measures (LME results: *= significant result, p < 0.05; ** = significant result, p < 0.01). Biparietal diameter (BPD), Occipitofrontal diameter (OFD), Head Circumference (HC), and Transcerebellar diameter (TCD).

| Volumetric/Linear measurements | DS /Controls |  | DS+CHD/DS-CHD |
| --- | --- | --- | --- |
|  | GA < 28.00 weeks | GA > 28.00 weeks |  |
| Whole brain and cortex |  |  |  |
| Supratentorial brain tissue (cm3) | 0.000** | 0.000** | 0.185 |
| Cortex (cm3) | 0.578 | 0.000** | 0.949 |
| BPD Skull (mm) | 0.731 | 0.464 | 0.069 |
| BPD Brain (mm) | 0.084 | 0.178 | 0.090 |
| OFD Skull (mm) | 0.142 | 0.000** | 0.233 |
| OFD Brain (mm) | 0.003** | 0.000** | 0.196 |
| HC (mm) | 0.578 | 0.003** | 0.107 |
| Cerebellum |  |  |  |
| Cerebellum (cm3) | 0.027* | 0.000** | 0.464 |
| TCD (mm) | 0.163 | 0.000** | 0.461 |
| Vermis height (mm) | 0.000** | 0.000** | 0.740 |
| Vermis width (mm) | 0.186 | 0.000** | 0.179 |
| Vermis surface area (mm2) | 0.000** | 0.000** | 0.722 |
| CSF regions |  |  |  |
| Lateral Ventricles (cm3) | 0.017* | 0.000** | 0.043* |
| eCSF (cm3) | 0.949 | 0.510 | 0.236 |
| Cavum width (mm) | 0.113 | 0.079 | 0.740 |

Fetuses and neonates with DS had significantly smaller brain occipitofrontal diameters (OFD) (p < 0.01, Fig 1f), in both the second and third trimester compared to age matched controls. Skull OFD (p < 0.01, Fig 1e) and HC (p < 0.01, Fig 1g) were only found to be significantly smaller in the third trimester, from 28 weeks GA (Table 3; Fig. 1). Skull and brain BPD was not significantly different between the two cohorts at any time point. There were no significant differences in 2D linear measurements between the DS fetuses with and without cardiac abnormalities (Table 3).

### Cerebellum

2D and volumetric cerebellar measurements in the DS cohort were significantly smaller than in the control population; cerebellar volume (p < 0.01), transcerebellar diameter (TCD) (p < 0.01), vermis height, width and surface area (p < 0.01) (Table 3; Fig. 2). Cerebellar volumes, vermis height and surface area were consistently smaller in the DS cohort, during the second trimester. The TCD and vermis width, however, were only smaller in the DS cohorts from the third trimester. Cerebellar volumes in late gestation DS fetuses and neonates were disproportionately smaller with respect to whole brain volumes, compared to the control population. No differences in cerebellar measures were found in the DS cohort with a CHD, compared to those without a CHD.

### Cerebrospinal Fluid (CSF) regions

Lateral ventricular volumes were larger in DS cases compared to controls throughout the second and third trimester (p < 0.01; Fig 3a). Lateral ventricular volumes were also found to be larger in DS cases without a CHD, compare to those with a CHD. Extra cerebral CSF (eCSF; Fig 3b) volumes and cavum pellucidum width were not significantly different (p > 0.05) (Table 3, Fig. 3).

## Discussion

Using state-of-the-art *in vivo* MR imaging, we have identified quantifiable alterations in early brain development in the fetus and neonate with DS, from as early as 21 weeks’ gestation. This study highlights growth trajectories of specific brain structures in the developing DS brain, and more importantly the varying time points during gestation when deviations arise. These include early second trimester deviations in whole brain and cerebellar volumes, and subsequent third trimester alterations in cortical and ventricular volumes, and with overall head circumferences. Despite the noted group-wise differences in brain development (DS compared to age-matched controls), we found overlap in almost all assessed parameters, highlighting a spectrum of atypical development across the DS cohort. This is of interest but perhaps not unexpected given the recognised individual variability in neurodevelopmental outcome of DS individuals, as established for instance, by IQ scores ^6,27–29^.

To the best of our knowledge this is the first study to use in-vivo MRI to detect early deviations in cerebellar and cortical growth during DS fetal development. Decreased cortical growth between 24 and 40 weeks gestational age is a likely biological substrate for impaired neurocognitive abilities in later childhood, especially complex functions relating to planning and attention ^30^. Reductions in brain size and altered cellular configurations, such as aberrant cortical lamination, reduction in dendritic ramifications and diminished synaptic formation ^31^ have all been detected post mortem in DS. More specifically human post-mortem morphometric and ultrastructural studies have shown a 20 - 50% reduction in neurons (from birth until 14 years of age), in the granular layers of the cerebral cortex along with an abnormal distribution of neurons in cortical layers II and IV ^32–34^. In addition studies that assessed dendritic morphology and arborisation in PM tissue have found that newborns and older infants with DS had shorter dendrites, a reduction in the number of spines and altered neuronal morphology (n=7; 26 weeks GA – 11 months) whilst DS fetuses exhibited the same neuronal morphology and dendritic spines as control fetuses (Takashima *et al.*, 1981). It is, therefore, likely that the anatomical volumetric reductions demonstrated with *in vivo* MRI reflect the changes in the underlying neurobiology. Decreasing growth trajectories over time, could be indicative of detrimental alterations in dendrite morphology and impaired cortical connectivity with increasing age. MRI provides an excellent and safe in-vivo assessment to monitor these changes across gestation and postnatally.

Our study also found reduced cerebellar volumes in the DS cohort, with more marked deviations from aged-matched controls arising with increasing PMA. This supports findings from previous MRI studies on adult DS populations where decreased cerebellar volumes were reported ^35–37^. Cerebellar volumetric abnormalities, have previously been associated with cognitive functions such as impaired attention, language, abstract reasoning, attention, working memory and executive control^38,39^. Neurodevelopmental assessments have demonstrated that children with DS performed worse than the typically developing children in the spatial-simultaneous working memory tasks, attributable to variations in cortical and cerebellar growth ^40^.

In addition, reductions in 2D linear measurements of vermian height, width and surface area were seen in the DS cerebellum compared with controls from 21 weeks GA. Whilst not conventionally performed in clinical practice, the fetal cerebellar vermis has successfully been measured on ultrasound ^41,42^ and it is possible that additional vermis height measures during the fetal anomaly US scan, might increase the non-invasive detection of fetuses with T21 at an earlier time point. Mouse models of DS have implicated the gene Dyrk1A in specific structural and functional cerebellar phenotypes, including reduced volume^43^. Cerebellar neurons are mainly formed prenatally, with some granule cell layer formation postnatally ^5^. Human post-mortem fetal studies have found evidence of impaired cellular formation, in the form of hypocellularity in the DS cerebellum which are the likely underlying cause of the gross volume reduction on MRI ^44,45^.

We have identified reduced 2D brain and skull parameters from as early as the second trimester, in addition to an altered ratio of OFD to BPD, which is consistent with previous reports of a brachycephalic skull in DS populations. The brachycephalic skull reflects a similarly shaped brain and is considered to be a consequence of relatively smaller frontal lobes in DS ^46^. Frontal lobe dysfunction and structural changes have previously been associated with early manifestations of Alzheimer’s disease in the adult DS population ^47,48^. Further analysis of regional brain volumes in early postnatal development will allow us to explore the relationship between frontal lobe development and childhood cognitive abilities.

The DS cohort had significant lateral ventricular enlargement at both the fetal and neonatal time points, compared to controls. However, eCSF volumes were similar between groups across gestation. Previous DS fetal ultrasound studies have reported enlargement of the lateral ventricles ^49^. Isolated fetal ventriculomegaly in non-DS children has been associated with mild cognitive impairment ^50^. Lateral ventricular dilation is known to increase across lifespan and is reflective of atrophy of surrounding brain tissue; this is particularly prominent in adults with AD pathology ^51^. Early ventricular enlargement is, however, likely to be developmental in origin. DS murine models have demonstrated that overexpression of the calmodulin regulator protein, Purkinje cell protein 4 (Pcp4), contributes to impaired ciliary function in ependymal cells and results in brain ventriculomegaly ^52^.

Whilst we hypothesised that regional brain volumes would be significantly smaller in the DS cohort with a cardiac defect, the only difference we identified antenatally was increased ventricular volume. Abnormalities in brain development of fetuses with CHD, without evidence of an underlying chromosomal abnormality, have reported increased incidences of ventriculomegaly ^53–55^. Other studies in neonates with congenital heart disease have focused on atypical growth and development of the cortex, which has been attributed to alterations in cerebral blood flow and oxygenation and the presence of transposition of the great arteries (TGA) and hypoplastic left heart ^56^. The cardiac conditions in our cohort with DS were primarily endocardial cushion defects such as AVSD, where antenatal changes in blood flow and oxygenation are less prominent than in TGA ^57^. Previous studies have reported a poorer outcome in individuals with DS who have an associated congenital heart defect (CHD) ^12–14^. More detailed studies of the postnatal brain development in DS with and without a cardiac defect may be enlightening, as it is possible that the specific genes responsible for a cardiac abnormality in DS may have an additional direct effect on brain development ^58^.

The primary limitation of this study is the small sample size of the DS cohort to date. DS pregnancy termination statistics were 90% in 2012, as reported by the National Down Syndrome Cytogenetic Register in England and Wales ^59^. The live-birth rate of DS has remained relatively stable over the last 30 years ^60^, consequently we found recruitment to be challenging. In addition, we combined images from a 1.5T and a 3T scanner for fetal imaging (all neonates were imaged on a 3T scanner). Images acquired from scanners with a stronger magnetic field and higher quality optimised receiver coils, benefit from an increased signal to noise ratio (SNR), consequently there may be differences in the acquired resolution and contrast of the resultant reconstruction. Previous studies found that while most of the brain was identically classified at the two field strengths, there were some regional differences observed ^61^. Others have reported highly reproducible volumetric data in adults with AD and controls, scanned on 1.5T and 3T scanners ^62^. Upon closer investigation when comparing datasets acquired on 1.5 T and 3T scanners, we were able to conclude that there were no discernible differences in volumetric data. Any differences fall within the accepted threshold of +/-5%.

In conclusion, this study has demonstrated alterations in both cortical and cerebellar development detectable by *in vivo* MR imaging *in utero* as early as the second trimester. Individuals with DS are now living to middle age, with improved quality of life and decreases in early mortality as a consequence of medical interventions, and improved surgical repair of congenital heart defects ^60,63,64^. The majority of individuals with DS, however, demonstrate cognitive decline in early adulthood. Strategies to modify this decline are already taking place ^65–67^ with researchers also testing therapeutic interventions to improve early brain development which may in turn decrease the likelihood of subsequent declines ^68^. Later neurodevelopmental assessments in our cohort will allow us to determine whether deviations in these antenatal indices are predictive of subsequent cognitive outcomes in DS and may help identify which children may need support, as early as possible.

## Acknowledgements

We thank the parents and children who participated in this study. The authors gratefully acknowledge staff from the Centre for the Developing Brain at King’s College London and the Neonatal Intensive Care Unit at St. Thomas’ Hospital. In particular, the research radiologists, radiographers, clinicians, neonatal nurses, midwives and the administrative teams. In addition, we wish to thank all our obstetric and fetal medicine colleagues from our patient identification sites who have referred participants to us.

This work was supported by the Medical Research Council [MR/K006355/1 and MR/LO11530/1]; Rosetrees Trust [A1563], Fondation Jérôme Lejeune [2017b – 1707], Sparks and Great Ormond Street Hospital Children’s Charity [V5318]. We also gratefully acknowledge financial support from the Wellcome/EPSRC Centre for Medical Engineering [WT 203148/Z/16/Z], the National Institute for Health Research (NIHR) Biomedical Research Centre (BRC) based at Guy’s and St Thomas’ NHS Foundation Trust and King’s College London and supported by the NIHR Clinical Research Facility (CRF) at Guy’s and St Thomas’. The Brain Imaging in Babies (BIBS) team additionally acknowledge support from EU-AIMS – a European Innovative Medicines Initiative; and infrastructure support from the National Institute for Health Research (NIHR) Mental Health Biomedical Research Centre (BRC) at South London and Maudsley NHS Foundation Trust and King’s College London.

The views expressed are those of the author(s) and not necessarily those of the NHS, the NIHR or the Department of Health.

