## Supplementary Materials for "Early alterations in cortical and cerebellar regional brain growth in Down Syndrome: An in-vivo fetal and neonatal MRI assessment"

| DS Subject ID | Fetal scan (GA weeks) | Neonatal scan (PMA weeks) | Sex (Male/ Female) | Cardiac abnormality | Other medical conditions |
|---|---|---|---|---|---|
| 101 | 21.86 | 42.71 | F | AVSD | x |
| 102 | 22.14 | | M | x | VM and agenesis of the corpus callosum |
| 103 | 22.29 | | M | AVSD | x |
| 104 | 25.29 | | F | AVSD | VM |
| 105 | 26 | | F | mild tricuspid regurgitation and mild right/left asymmetry | x |
| 106 | 26.29 | | M | coarctation of the aorta | x |
| 107 | 26.57 | | M | cardiac asymmetry | x |
| 108 | 27.14 | | F | x | VM |
| 109 | 27.29 | | M | Tetralogy of Fallot | VM and duodenal atresia |
| 112 | 28.14 | | M | x | VM |
| 113 | 29.43 | | M | x | agenesis of cerebellar vermis |
| 114 | 29.86<br>35.71 | 39.71 | M | AVSD and hypoplastic arch | x |
| 115 | 31 | 43.00 | F | balanced AVSD and right sided aortic arch | x |
| 116 | 31 | | F | x | Twin case – baby small for GA. Other twin is a normally developing baby |
| 117 | 33.29 | | M | x | VM and oesophageal atresia |
| 118 | 33.29 | 41.29 | F | VSD | x |
| 119 | 33.57 | | F | x | VM |
| 120 | 33.86 | | M | VSD | x |
| 121 | 34.43 | 44.57 | M | x | x |
| 122 | 34.86 | | M | Tetralogy of Fallot with severe pulmonary stenosis | x |
| 123 | 35 | | F | AVSD | x |
| 124 | 35.43 | 39.00 | M | x | duodenal atresia |
| 201 | | 41.86 | M | AVSD | x |
| 202 | | 43.43 | F | x | x |
| 203 | | 40.57 | F | x | duodenal atresia |
| 204 | | 36.14 | F | VSD and pulmonary stenosis | duodenal atresia |
| 205 | | 41.43 | F | AVSD and PDA | duodenal atresia |
| 206 | | 43.00 | M | ASD | Hirschsprung’s |
| 207 | | 45.57 | F | Tetralogy of Fallot | Hirschsprung’s |
| 208 | | 38.43 | F | Tetralogy of Fallot, PDA | x |
| 209 | | 43.57 | F | AVSD and hypoplastic arch | x |
| 210 | | 44.57 | M | x | x |
| 211 | | 44.71 | F | VSD with bidirectional shunt and large PDA | Congenital hypothyroidism |

| | | | | | |
|---|---|---|---|---|---|
| 212 | | 32.43 | F | mildly dysplastic aortic valve | x |
| 213 | | 37.57 | M | AVSD and coarctation of the aorta | x |
| 214 | | 38.86 | M | ASD | Hirschsprung’s |
| 215 | | 37.86 | F | AVSD | x |
| 125 | 30.57 | | F | AVSD | - |
| 126 | 31.43 | | M | x | x |
| 127 | 34.14 | | M | x | x |
| 128 | 30.14 | | F | x | x |
| 129 | 30.43 | | M | x | x |
| 130 | 30.71 | | F | right sided aortic arch | x |
| 131 | 27.29 | | F | aberrant right subclavian artery and intermittent tricuspid regurgitation | VM |

Table 4: Age at scan and clinical characteristics of fetuses and neonates with DS. Atrioventricular septal defect (AVSD), atrial septal defect (ASD), ventricular septal defect (VSD), patent ductus arteriosus (PDA) and ventriculomegaly (VM)
